# Bidirectional Control of Coronary Vascular Resistance by Eicosanoids via a Novel GPCR

**DOI:** 10.1101/420406

**Authors:** Nabil J. Alkayed, Zhiping Cao, Zu Yuan Qian, Shanthi Nagarajan, Xuehong Liu, Jonathan Nelson, Fuchun Xie, Bingbing Li, Wei Fan, Lijuan Liu, Marjorie R. Grafe, Xiangshu Xiao, Anthony P. Barnes, Sanjiv Kaul

**Affiliations:** Department of Anesthesiology & Perioperative Medicine, Oregon Health & Science University, Portland.; Department of Physiology & Pharmacology, Oregon Health & Science University, Portland.; Department of Pathology, Oregon Health & Science University, Portland.; The Knight Cardiovascular Institute, Oregon Health & Science University, Portland.

**Author notes:** Correspondence to: S.K., A.P.B., X.X. or N.J.A.

## Abstract

Arachidonic acid metabolites epoxyeicosatrienoates (EETs) and hydroxyeicosatetraenoates (HETEs) are important regulators of myocardial blood flow and coronary vascular resistance (CVR), but their mechanisms of action are not fully understood. We identified G protein-coupled receptor 39 (*GPR39)* as a microvascular smooth muscle cell (mVSMC) receptor antagonistically regulated by two endogenous eicosanoids: 15-HETE, which stimulates GPR39 to increase mVSMC intracellular calcium and augment microvascular CVR, and 14,15-EET, which inhibits these actions. Furthermore, zinc ion acts as an allosteric modulator of GPR39 to potentiate the efficacy of the two ligands. Our findings will have a major impact on understanding the roles of eicosanoids in cardiovascular physiology and disease, and provide an opportunity for the development of novel GPR39-targeting therapies for cardiovascular disease.

**One Sentence Summary:** GPR39 is a microvascular smooth muscle cell receptor regulated by two vasoactive eicosanoids with opposing actions.

---

Myocardial oxygen demand determines coronary blood flow, which is exquisitely regulated through changing vascular resistance in coronary arterioles ranging in size from 100µm to 300µm (*1*). Whereas the determinants of myocardial oxygen demand are well known, the molecules responsible for the second-to-second regulation of microvascular tone and the receptors through which they act have not been identified. Two classes of signaling P450 eicosanoids, epoxyeicosatrienoates (EETs) and hydroxyeicosatetraenoates (HETEs), are potent regulators of microvascular tone and play critical roles in cardiovascular physiology and disease (*2, 3*). EETs are predominantly microvascular dilators and have been shown to function as endothelium-derived hyperpolarization factors (EDHFs) in the coronary circulation (*4, 5*), and to mediate functional hyperemia in the brain (*6, 7*). They are anti-inflammatory, inhibit platelet aggregation and protect against ischemia-reperfusion injury in heart and brain (*8*). HETEs, on the other hand, are typically vasoconstrictors that have been implicated in the generation of arteriolar myogenic tone and blood flow autoregulation in the brain and kidneys (*9, 10*). They promote inflammation and contribute to hypertension and ischemia-reperfusion injury *(11)*.

The mechanisms utilized by these eicosanoids to exhibit their biological actions in microvascular smooth muscle cells (mVSMCs) remain largely unknown. Ligand binding and pharmacological studies suggest that EETs signal via a G protein-coupled receptor (GPCR) *(12)*, yet the identity of an EETs receptor has remained elusive. In the current study, we generated a clickable photocrosslinking probe based on 14,15-EET that allowed us to isolate its binding proteins from mouse heart mVSMCs in an unbiased manner using mass spectrometry-based proteomics. This approach identified GPR39, a member of the Ghrelin peptide receptor family, as a putative receptor for 14,15-EET in mVSMCs. Here, we show that GPR39 is capable of mobilizing calcium in VSMC in response to 15-HETE as well as serving as the site of 14,15-EET’s inhibitory action on 15-HETE-induced calcium signaling, indicating that it acts as a receptor for both eicosanoids. We also localize GPR39 immunoreactivity to mouse heart mVSMCs which is consistent with a role in microvascular tone regulation. Finally, we demonstrate that 15-HETE increases CVR using the isolated Langendorff mouse heart perfusion preparation, an effect that is inhibited by co-administration of 14,15-EET and requires GPR39 expression.

## Results

### Identification of GPR39 as a putative receptor for 14,15-EET

We implemented a chemical biology approach to purify 14,15-EET-binding proteins by producing a modified form of 14,15-EET (EET-P, **Fig. 1A**) that covalently crosslinks to target proteins following exposure to ultraviolet (UV) light. The probe also incorporates a click chemistry moiety that allows subsequent fluorophore labeling or biotin-streptavidin affinity purification and identification of linked proteins by mass spectrometry (**Fig. 1A**). Despite these additional functional groups, we were able to demonstrate that EET-P mimics 14,15-EET’s previously reported ability to dilate mouse mesenteric arteries pre-constricted with thromboxane A2 agonist U46619 (*13*). We next used this probe to identify putative mVSMC 14,15-EET receptor(s) by treating mouse heart mVSMCs with EET-P (1 μM) in the presence or absence of 14,15-EET followed by a 5 min UV (365 nm) exposure. Cell lysates were reacted with biotin-azide for affinity purification and subsequent mass spectrometry analysis that yielded a number of intracellular and membrane-associated proteins that could be competitively displaced by 14,15-EET including a known 14,15-EET metabolizing enzyme, epoxide hydrolase, an indication of probe specificity. A single GPCR was detected in the screen, GPR39, a 50 kDa orphan member of the ghrelin receptor family previously reported to be activated by zinc ions (Zn^2+^) (*14*). Rhodamine-azide labeling followed by SDS-PAGE analysis detects an EET-P-labeled protein that was dose-dependently displaced by 14,15-EET and approximately the same molecular weight as GPR39 (**Fig 1B**). Similarly, we find that cultured mVSMCs can be labeled with EET-P and an in-cell click reaction with a rhodamine azide. Importantly, this labeling can be displaced by pretreatment with 14,15-EET (**Fig. 1C**). The GPR39 genomic locus encodes two isoforms, GPR39 1a (a full-length 7-transmembrane (7TM) isoform) and 1b (a truncated 5TM 1b isoform that lacks TM6 and 7) (*15*). Using immunocytochemistry and real-time quantitative PCR we confirmed expression of GPR39 1a, but not 1b in cultured mouse heart mVSMCs (**Fig. 1D**). We also determined that human embryonic kidney (HEK)-293 cells express only the 1b isoform. Probe binding specificity was confirmed by crosslinking EET-P to HEK-293 cells transiently transfected with HA-tagged human GPR39 1a. Western blots probed with HA-antisera (**Fig. 1E**) indicate that HA-tagged GRP39 1a co-purifies with EET-P (lane 1) and this band is absent in lysates of untransfected cells (lane 4). We further confirmed probe specificity by eliminating probe binding to HA-tagged GPR39 by pretreating GPR39-transfected HEK-293 cells with either 14,15-EET (5 μM, lane 2) or EETs antagonist 14,15-epoxyeicosa-5(Z)-enoic acid (*16*) (14,15-EEZE, 5 μM, lane 3) prior to EET-P exposure. Finally, dot-blot assay demonstrated dose-dependent saturable binding of 14,15-EET, but not 11,12-EET, to protein extracts from HEK-293 cells expressing GPR39 1a, but not control cells (**Fig 1F**; raw images above, and optical density quantification below).

**Fig. 1.**
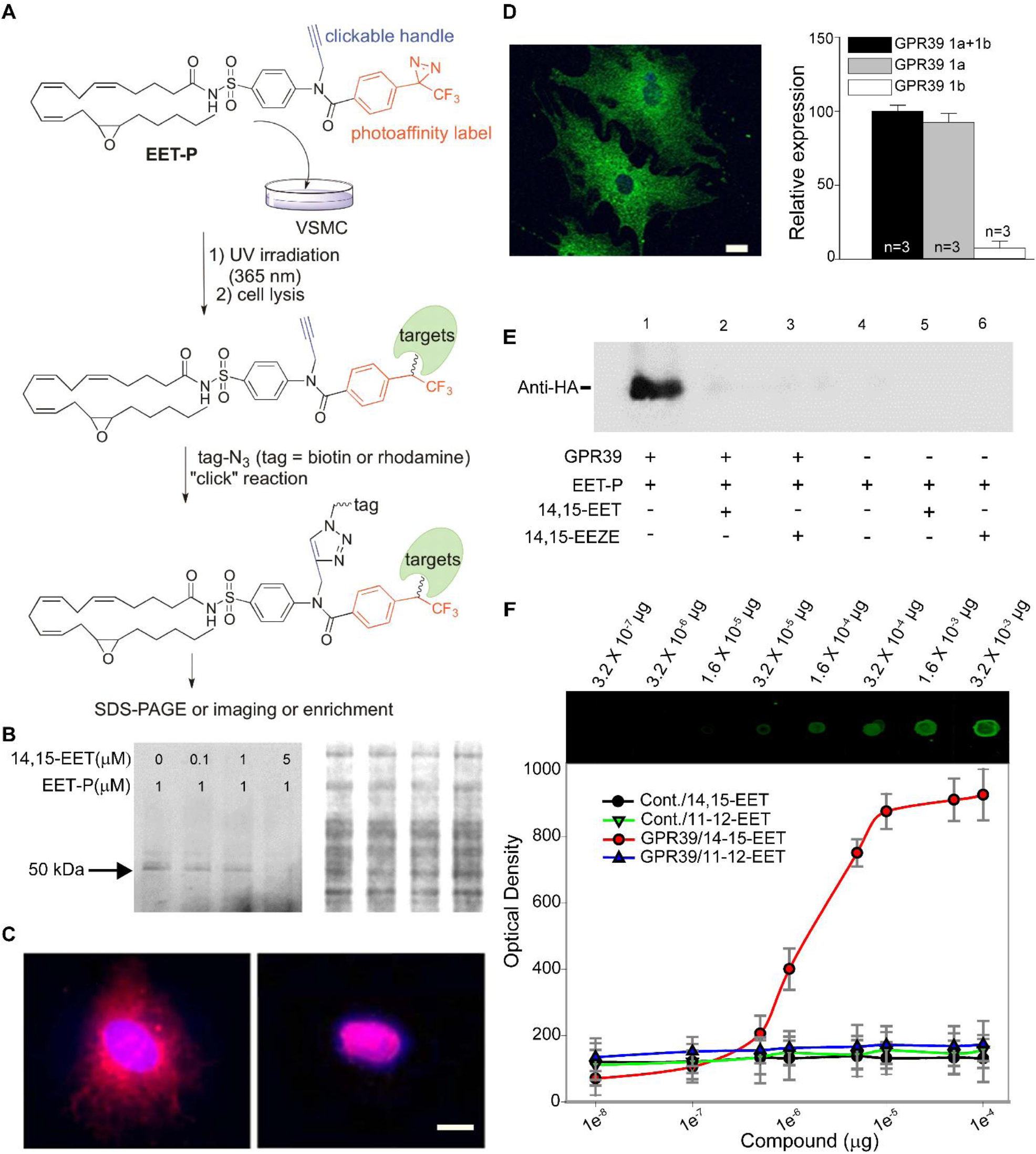
Purification and validation of GPR39 as a putative receptor for 14,15-EET. (**A)** Chemoproteomics strategy to identify 14,15-EET receptor in mVSMCs. (**B**) Photocrosslinking GPR39 with EET-P in mVSMCs. Left: SDS-PAGE gel revealed the presence of a ∼50 kDa band. 14,15-EET pre-treatment reduced protein binding to EET-P in a dose-dependent manner. Right: Total protein determined by Coomassie Blue staining of the same gel. (**C**) Confocal images demonstrating binding of EET-P to outer surface of mVSMCs (left, red). Pretreatment with 1 μM 14,15-EET for 10 min prevented EET-P surface binding (right). (**D**) A representative immunofluorescent confocal image illustrating GPR39 expression (left, green) in primary cultured mouse heart mVSMCs (scale bar = 10 μm), and expression of GPR39 1a, but not 1b, confirmed by qPCR (right). (**E**) Western blot detection of EET-P crosslinking to epitope-tagged GPR39 1a in transfected HEK cells. HEK cells were treated with 5 μM 14,15-EET or 14,15-EEZE for 10 min, 1 μM EET-P for 15 min, and then irradiated with UV for 5 min at 4^°^C. After clicking with biotin and purification with streptavidin Dynabeads, protein extracts were probed by anti-HA antibody. (**F**) Dot-blot assay illustrating dose-dependent binding of 14,15-EET, but not 11,12-EET, to membrane protein extracts from HEK-293 cells stably expressing GPR39 1a, but not untransfected control cells (raw images on top, and optical density quantification at the bottom).

### Isoform-Specific Activation of GPR39 Signaling by 14,15-EET and 15-HETE

We transfected HEK-293 cells with either GPR39 1a or GPR39 1b to determine if 14,15-EET binding activates GPR39 signaling and if 14,15-EET binding is specific or whether related eicosanoids can also activate this receptor. Cells were subsequently stimulated with one of four regioisomers of EETs (5,6-EET; 8,9-EET; 11,12-EET; 14,15-EET), HETEs (11-HETE, 12-HETE, 15-HETE, 20-HETE), or vehicle. 14,15-EET and 15-HETE are the only regioisomers that significantly increase GPCR activation as monitored by ERK phosphorylation in cells transfected with GPR39 1a (**Fig. 2A**, **2B**) but not GPR39 1b or untransfected cells (**Fig. 2C-D**), with both eicosanoids displaying a concentration-dependent activation of GPR39 (**Fig. 2C** and **2D)**. Dot-blot assays using lysates from GPR39 1a stably expressing HEK-293 cells demonstrate dose-dependent, saturable binding of 15-HETE, similar to 14,15-EET, that is not observed for 12-HETE. Importantly, binding competition experiments indicate that 15-HETE and 14,15-EET can displace each other while 12-HETE and 11,12-EET, respectively, fail to do so.

**Fig. 2.**
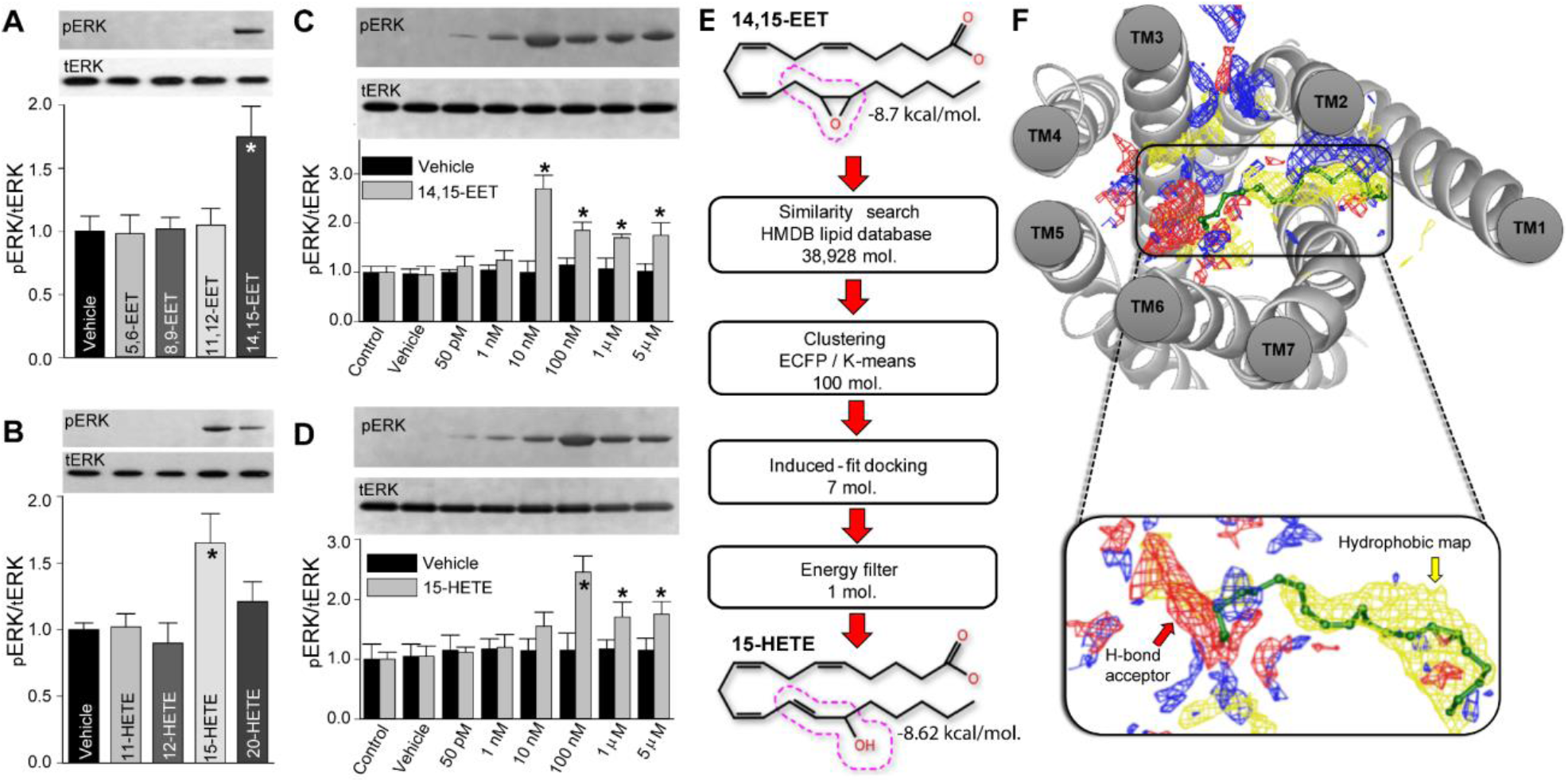
Functional activation and modeling of GPR39 ligand binding. ERK phosphorylation induced by 1 min treatment with 1 μM of each one of 4 regioisomers of EETs (**A**) or HETEs (**B**). Total ERK was used as a protein loading control. Dose-dependent ERK phosphorylation induced by 1 min treatment with either 14,15-EET (**C**) or 15-HETE (**D**) in HEK cells expressing GPR39 1a, but not untransfected cells (ANOVA, n=6/treatment, p-value <0.005). (**E**) Virtual Screening of 14,15-EET-like compounds from HMDB database revealed comparable energies for 15-HETE and 14,15-EET. The 14,15-EET and 15-HETE structures exhibit a high degree of structural similarity. The dotted red line around carbons 13-15 highlights the structural difference between the two eicosanoids. (**F**) GPR39 7TM core binding pocket and an enlarged view of the predicted binding pose of 14,15-EET. Blue, red and yellow maps correspond to hydrogen bond donor, hydrogen bond acceptor and hydrophobic areas within the binding pocket, respectively. N-terminal residues contributing to minor pocket are not shown for the sake of clarity. The carboxylate group of 14,15-EET interacts with polar residues in TM6 and the lipid portion interacts with the hydrophobic site formed at the minor-binding pocket, as shown in site mapping.

### GPR39-Ligand Interaction Modeling Predicts Selectivity for 14,15-EET and 15-HETE

We next used a homology model to investigate GPR39 1a-ligand specificity in silico that predicted structural complementarity between 14,15-EET and GPR39, as indicated by a low docking energy (−8.7 kcal/mol). A fingerprint similarity search for other GPR39 ligands using 14,15-EET as a reference structure resulted in a list of long-chain fatty acids including 15-HETE, which had a predicted docking energy of −8.6 kcal/mol, similar to 14,15-EET. This likely reflects that both compounds have a polar group at the 15^th^ carbon position and maintain double bonds in the *cis* configuration at the 5^th^, 8^th^ and 11^th^ positions (**Fig. 2E**) unlike the other EETs and HETEs regioisomers. A potential orthostatic pocket was identified using SiteMap V3.2 which revealed a hydrophilic major pocket and a hydrophobic minor pocket in GPR39 1a with the tail portion of these lipids extending into the minor binding pocket formed by transmembrane 1 (TM1), TM2, TM7 and the N-terminal loop (**Fig. 2F**). The major pocket accommodates the carboxylate moiety of 14,15-EET and 15-HETE by forming ionic interactions with positively charged residues from TM6 (**Fig. 2F**). These structural predictions and their validation using heterologous expression strongly support activation of GRP39 by 15-HETE.

### GPR39 Mediates Effects of 14,15-EET and 15-HETE on mVSMC Calcium Transients

We next enquired whether 14,15-EET and 15-HETE regulate the signaling of mVSMC via GPR39. Using live-cell fluorescent imaging to monitor calcium transients, we found that 15-HETE increased intracellular calcium in mVSMCs (**Fig. 3A**), and GPR39 RNAi knockdown in mVSMCs attenuated the increase in mVSMC intracellular calcium produced by 15-HETE (**Fig 3B**), confirming a significant role for GPR39 in mediating the effect of 15-HETE on mVSMC calcium transients. 14,15-EET had no effect on intracellular calcium in mVSMCs at concentrations between 1 pM - 10 μM (**data not shown**), but it was observed to inhibit 15-HETE-dependent increases in intracellular calcium at 1 μM (**Fig. 3C,D**), consistent with GPR39 acting as a dual sensor for both 14,15-EET and 15-HETE. Taken together, our modeling data along with pERK, calcium imaging and binding assays indicate a competitive structure-activity relationship between 14,15-EET and 15-HETE.

**Fig. 3.**
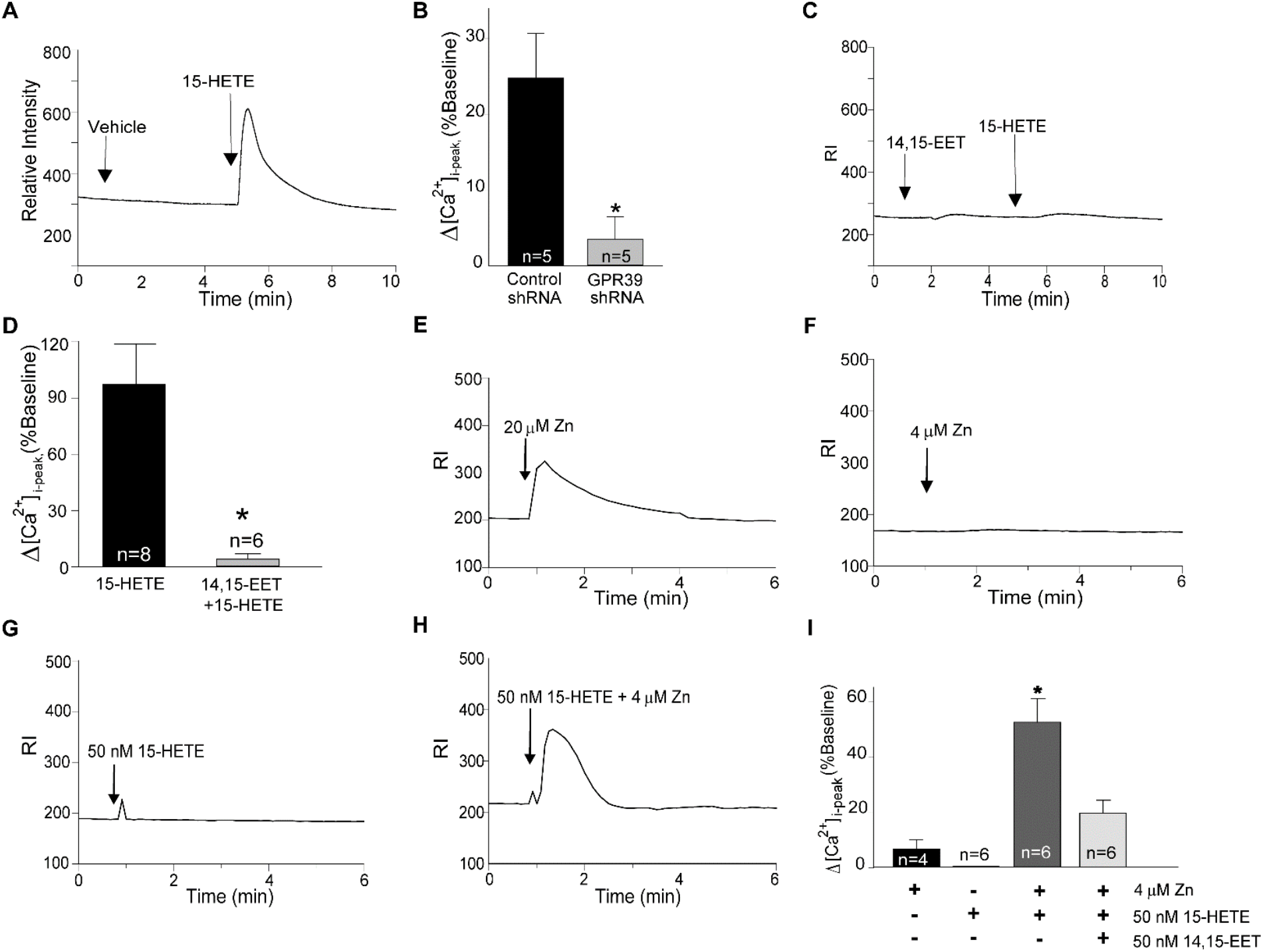
**(A)** Calcium imaging of mVSMCs showing an increase in mVSMCs [Ca^2+^]i by 1 μM 15-HETE. **(B)** Summary of [Ca^2+^]i response to 1 μM 15-HETE in mVSMCs treated with a lentivirus containing either a scrambled or GPR39-targetting shRNA for 72 hours. **(C, D)** The increase in mVSMCs [Ca^2+^]i by 15-HETE is abolished by pre-treatment with 1 μM 14,15-EET. (**E**) Zinc alone increases [Ca^2+^]i at concentrations above 20 μM, but neither zinc at 4 μM **(F)** nor 15-HETE at 50 nM **(G)** alone had an effect on [Ca2+]i. **(H)** Zinc (4 μM) potentiates the effect of 15-HETE (50 nM) on [Ca^2+^]i in mVSMCs. **(I)** Summary of changes in mVSMCs [Ca^2+^]i in response to 50 nM of 14,15-EET and 15-HETE, separately and in combination, and with and without zinc.

### Zinc Sensitizes GPR39 to 14,15-EET and 15-HETE

Zinc has been reported to be either a GPR39 agonist (*14*) or an allosteric modulator for synthetic ligands of the receptor (*17*). Therefore, we first tested the effect of Zn^2+^ alone in mVSMCs and found that it increased intracellular calcium at concentrations ≥20 μM **(Fig 3E**) and had no effect at concentrations between 1-10 μM (**Fig 3F**). At these lower concentrations, however, we observe it potentiating the response of GPR39 to 15-HETE stimulation, consistent with a role as an allosteric modulator. Application of 15-HETE alone elicits a calcium response in mVSMCs only at concentrations at or above 100 nM (**Fig. 3G**). However, in the presence of 4 µM Zn^2+^, 15-HETE elicits a significant calcium response in mVSMCs at a concentration of 50 nM (**Fig. 3H and I**), which was inhibited by 50 nM 14,15-EET (**Fig 3I**). Overall, these results suggest that zinc serves as a positive allosteric modulator for these endogenous ligands.

### GPR39 Localization and Function in Myocardial Microvessels

Consistent with a role in regulating coronary vascular resistance, immunofluorescence indicates that GPR39 expression is predominantly restricted to *microvessels* in the mouse heart (**Fig 4A-D)**. Co-staining with α-smooth muscle actin (α-SMA) confirms expression in arteriolar smooth muscle cells (**Fig 4B, C**). Vascular smooth muscle localization was further confirmed by co-localization with red fluorescence in hearts from the NG2-DsRed transgenic mouse (**Fig 4D**). This microvascular pattern of expression was further confirmed using non-fluorescent immunostaining where we also observed GPR39 expression in both mouse and human heart within perivascular cells that did not label with α-SMA but were consistently adjacent to microvessels including capillaries, suggestive of GPR39 expression in pericytes as well.

**Fig. 4.**
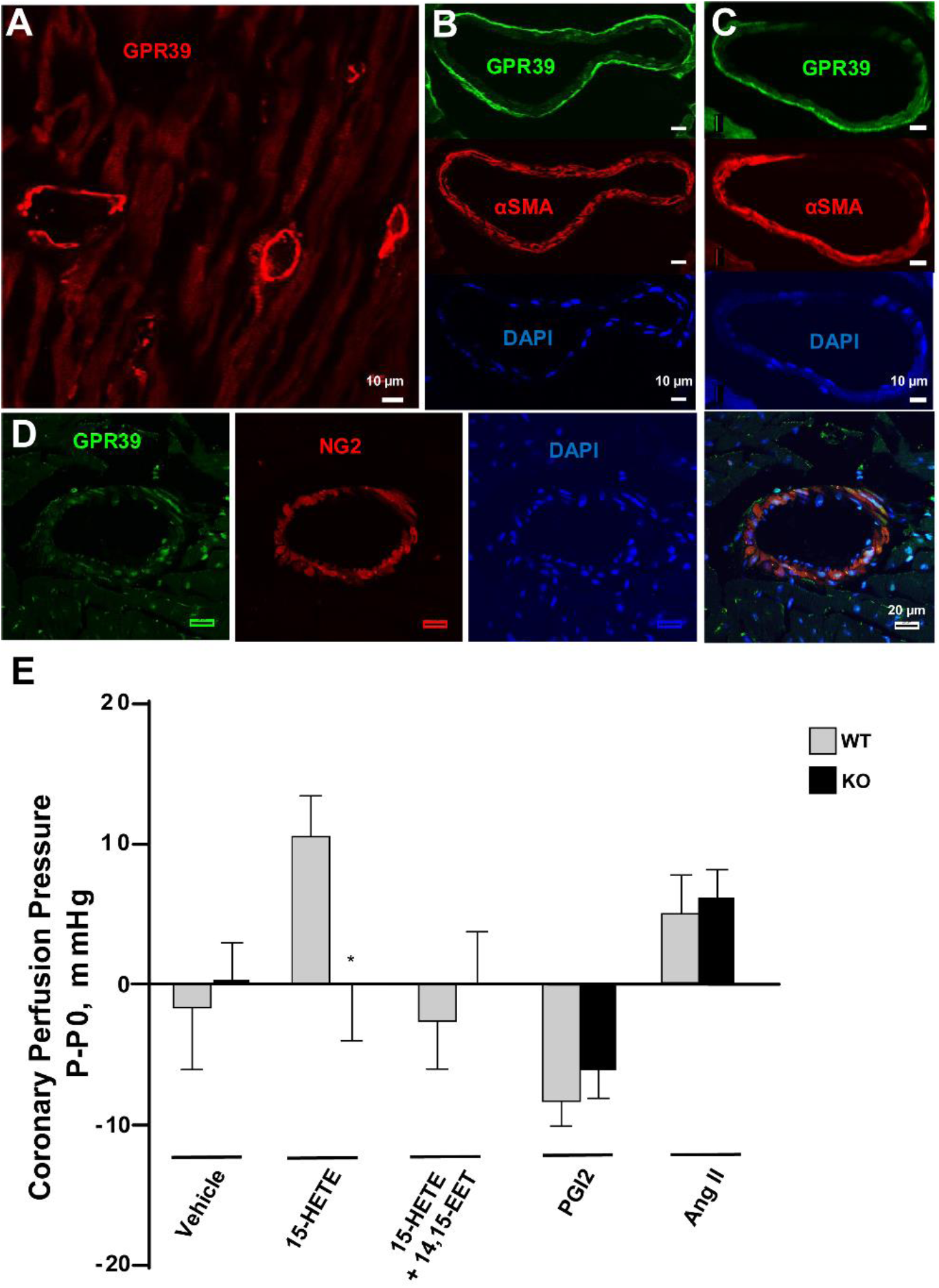
Microvascular localization and function of GPR39 in mouse heart. (**A-D**) Immunofluorescent imaging of GPR39 shows microvascular pattern of expression in mouse heart (upper left, red; scale bar = 10 μm), which co-localizes with VSMC markers *α*-smooth muscle actin (*α*-SMA, red, upper middle and right; scale bar = 10 μm) and NG2 (red, lower panel; scale bar = 20 μm). DAPI stains nuclei in blue. (**E**) Changes in coronary perfusion pressure, at a constant flow rate, in response to infusion of 15-HETE (1 μM), 15-HETE plus 14,15-EET (1 μM), prostaglandin I_2_ (PGI_2_, 200 nM), angiotensin II (AngII, 100 nM) or vehicle in isolated mouse heart preparation from wild-type (WT) and GPR39 knockout (KO) mice.

We next determined the contribution of GPR39 activation by eicosanoids to microvascular tone regulation. This was accomplished using a mouse heart Langendorff preparation where changes in coronary perfusion pressure (CPP) are determined by microvascular resistance at a constant flow rate. Consistent with its established role as a microvascular vasoconstrictor, 15-HETE (1 µM) increases CPP **(Fig 4E)**, and this effect is inhibited by co-administration of 14,15-EET (1 µM) or abolished in GPR39-null hearts (**Fig 4E**). Importantly, the effects of two other vasoactive agents on CPP, angiotensin II (Ang II) and prostaglandin I_2_ (PGI_2_), were unaffected by GPR39 deletion. These results support the notion that GRP39 is responsible for microvascular tone regulation by 15-HETE and 14,15-EET.

## Discussion

We developed a chemical crosslinking mimetic to identify 14,15-EET as an endogenous ligand of GPR39, canonically considered to be a zinc receptor. We further show that 14,15-EET and 15-HETE can regulate mVSMC calcium dynamics via their opposing actions on GPR39. We also provide evidence that the bidirectional control of coronary vascular resistance by these eicosanoids is dependent on the presence of GPR39.

Models of 14,15-EET signaling mediated by a GPCR have been proposed by several groups (*13, 18, 19*), yet the receptor has remained unidentified. Photoaffinity labeling was previously used to demonstrate a ∼47 kDa high-affinity binding protein of unknown identity for 14,15-EET in membrane fractions from U937 and vascular cells *(13)*. Unlike previous approaches, our clickable photocrosslinking probe allowed enrichment of crosslinked proteins from mVSMCs, permitting target identification by mass spectrometry. GPR39 represents the first high-affinity receptor for any EET regioisomer, although several membrane proteins have been shown to respond to high micromolar concentrations of 14,15-EET including: large conductance calcium-activated (BK_Ca_) and ATP-sensitive potassium channels, the transient receptor potential cation channel subfamily V member 4 (TRPV4), and some GPCRs (*13*), including GPR40 *(20)* and the prostaglandin E (EP2) *(18)* and thromboxane (TP) *19* receptors, as well as our own recent work screening a variety of GPCRs for 14,15-EET responses (*21*). Our modeling predictions also led us to test and confirm that 15-HETE is an additional endogenous ligand of GPR39. No high affinity receptor has been identified for 15-HETE, although receptors have been identified for other HETE regioisomers, including 12-HETE (*22*) and 20-HETE (*23*).

Endogenous ligands for GPR39 have remained elusive as early reports suggested obestatin, a ghrelin-derived peptide, but this has since been refuted *(14, 24)*. Multiple studies have linked GPR39 with metabolism including two studies of GPR39 knockout mice, one where body weight and food intake were reported to be normal *(25)*, while the other noted higher body weight and fat composition with no change in food intake (*26*). A report has also linked GPR39 loss with impaired insulin secretion (*27*), yet synthetic agonists of GPR39 fail to drive insulin secretion (*28*), and still others associate GPR39 loss with nervous system symptoms such as seizures *29* and depression-like behavior (*30*). GPR39 has also been proposed to function as a Zn^2+^ receptor *(14)*. However, it is unclear if the pharmacokinetics of Zn^2+^ with GPR39 support it as a physiological agonist of the receptor. It is more likely that zinc modulates receptor activity based on studies characterizing synthetic GPR39 agonists (*17*). Furthermore, Zn^2+^ is known to be an allosteric modulator for multiple receptors and ion channels (*31*). Here, we confirm GPR39 sensitivity to high concentrations of Zn^2+^, and critically, provide a novel function for physiologic levels of Zn^2+^ as an allosteric modulator enhancing the efficacy of the natural GPR39 ligands 14,15-EET and 15-HETE. Thus, Zn^2+^ may also play a role in the regulation and disorders of the microcirculation through GPR39 modulation. Recent reports suggested that zinc regulates endothelial function (*32*) and inhibits phosphate-induced vascular calcification (*33*), presumably through its action on GPR39.

In conclusion, our study is the first to propose a microvascular role for GPR39, and furthermore that this receptor senses the relative concentrations of 14,15-EET and 15-HETE in order to regulate mVSMC tone. It is very possible that an imbalance between these two eicosanoids could alter GRP39 activity, predisposing individuals for microvascular disease and microvascular complications of systemic diseases such as diabetes, hypertension and septic shock. Work by numerous investigators over the past three decades has established a critical role for P450 eicosanoids in cardiovascular physiology and disease (*34-39*). Further progress in this field has been hampered by the lack of understanding of the molecular mechanisms of actions of these important lipid mediators. Our identification of the receptor for two important endogenous vasoactive eicosanoids and their unique mode of dual and opposing regulation of receptor function is a major breakthrough in the field. These findings will have a significant impact on our understanding of how P450 eicosanoids control the microcirculation, their roles in microvascular disease, and the development of novel therapeutic agents for vascular disease.

## Acknowledgments

Mass spectrometric analysis was performed by the OHSU Proteomics Shared Resource with partial support from NIH core grants P30EY010572, P30CA069533, and shared instrumentation grant S10OD012246.

